# Divergence of *germ cell-less* roles in germ line development across insect species

**DOI:** 10.1101/2025.03.26.645619

**Authors:** Jonchee A. Kao, Ben Ewen-Campen, Cassandra G. Extavour

**Affiliations:** Department of Molecular and Cellular Biology, Harvard University, 16 Divinity Ave, Cambridge, MA 02138, USA; Department of Organismic and Evolutionary Biology, Harvard University, 16 Divinity Ave, Cambridge, MA 02138, USA; Department of Genetics, Blavatnik Institute, Harvard Medical School, Harvard University, Boston, MA 02115, USA; Howard Hughes Medical Institute, Chevy Chase, MD, USA

**Author notes:** Corresponding author: Cassandra G. Extavour.

**Keywords:** Germ cells, pole cells, *Oncopeltus fasciatus*, *Gryllus bimaculatus*, Hemimetabola, milkweed bug, cricket, Torso

## Abstract

During development, sexually reproducing animals must specify and maintain the germ line, the lineage of cells that gives rise to the next generation of animals. In the fruit fly *Drosophila melanogaster*, *germ cell-less* (*gcl*) is required for the formation of primordial germ cells in the form of cells that cellularize at the posterior pole of the embryo, called pole cells. Forming pole cells is a mechanism of germ cell formation unique to a subset of insects. Even though most animals do not form pole cells as primordial germ cells, *gcl* is conserved across Metazoa, raising the question of how this conserved gene acquired its central role in the evolutionarily derived process of pole cell formation. Here, we examine the functions of *gcl* in two other insects with different modes of germ cell specification: the milkweed bug *Oncopeltus fasciatus* and the cricket *Gryllus bimaculatus*. We found that *gcl* is involved in germ cell development, but not strictly required for germ cell specification, in *O. fasciatus*, although it appears to function through a mechanism different from that in *D. melanogaster*. In contrast, we could not detect any impact on the embryonic germ line upon *gcl* knockdown in *G. bimaculatus*. This work serves as a case study into how the roles of genes in the process of germ line development can change over evolutionary time across animals.

**Highlights:**

- *germ cell-less* knockout reduces germ cell number in *Oncopeltus fasciatus*.
- *germ cell-less* knockdown does not affect germ cell number in *Gryllus bimaculatus*.
- The genetic mechanisms through which *gcl* affects germ cell number have diverged among insects.

**Graphical abstract:** 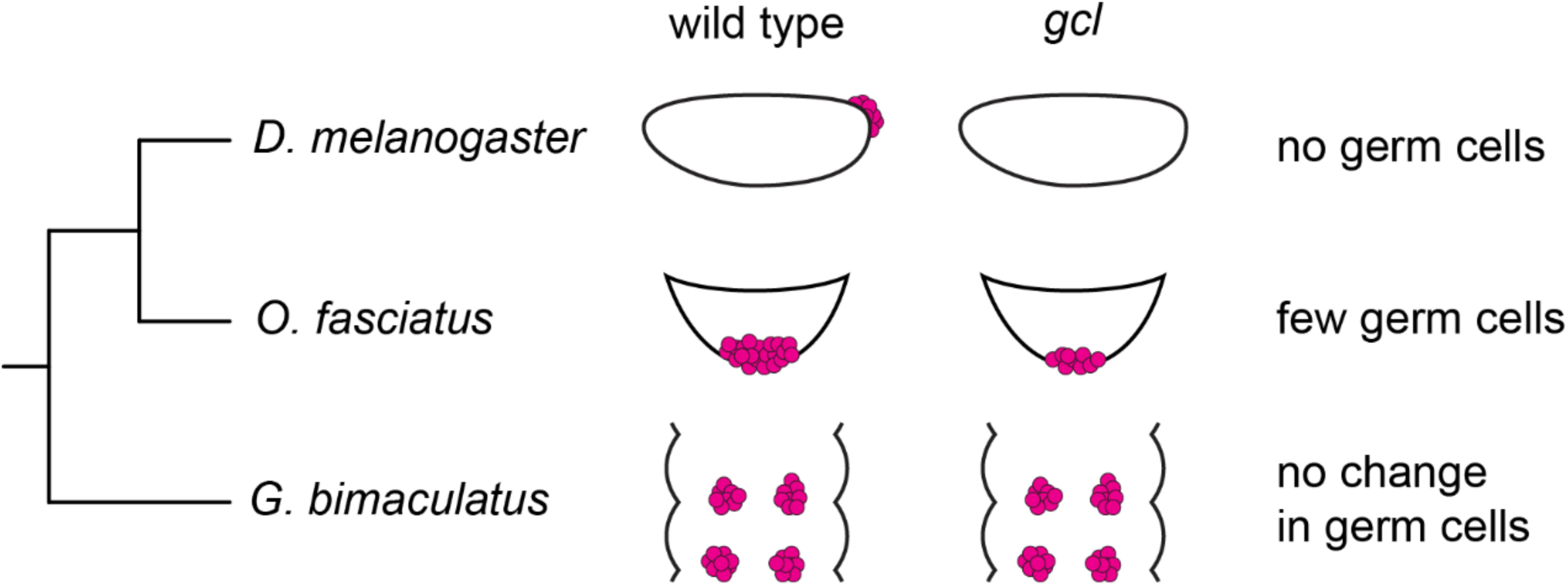

## Introduction

One of the most crucial processes in animal development is the specification and maintenance of the germ line, the lineage of cells uniquely capable of giving rise to the next generation of animals (Extavour and Akam, 2003). Despite the importance of this process, mechanisms of germ line development vary widely even within closely related taxa, including across insects (Extavour and Akam, 2003). For example, in the fruit fly *Drosophila melanogaster*, germ cells first appear at the posterior pole of the embryo very early in embryogenesis, before cellularization occurs in the rest of the embryo (Huettner, 1923). The formation of germ cells in embryos of this fly depends on the presence of the germ plasm, a specialized region of the cytoplasm at the posterior pole containing maternally synthesized RNAs and proteins that collectively direct the formation and specification of germ cells (Illmensee and Mahowald, 1974). In contrast, germ cells in the cricket *Gryllus bimaculatus* are first detectable comparatively later in embryogenesis, after segmentation of the abdomen has begun (Ewen-Campen et al., 2013a). There is no evidence of germ plasm in cricket embryos, and germ cells are thought to be specified instead by inductive signals from neighboring cells, including BMP signaling (Donoughe et al., 2014; Ewen-Campen et al., 2013a). In yet another mechanism of insect germ cell specification, germ cells of the milkweed bug *Oncopeltus fasciatus* appear at a stage intermediate between those of *D. melanogaster* and *G. bimaculatus*, after blastoderm cellularization but before abdominal segmentation (Butt, 1949; Ewen-Campen et al., 2013b). There is also no evidence of germ plasm in embryos of this bug (Ewen-Campen et al., 2013b), but specific germ cell specification signals have not been identified.

Despite the variation in modes of germ line development across species, many genes involved in this process are conserved. These include *vasa* and *piwi*, which are commonly used as germ cell markers in many different animals (Ewen-Campen et al., 2010). *germ cell-less* (*gcl*) is also sometimes used as a germ cell marker, but its role in germ line development is less well understood (Jongens et al., 1992). Embryos laid by *D. melanogaster* females homozygous for a loss of function mutation in *gcl* have significantly fewer germ cells than controls, if any (Jongens et al., 1994, 1992). The gene is thought to have a role in suppressing somatic cell fate by targeting the receptor tyrosine kinase Torso for degradation (Colonnetta et al., 2021; Pae et al., 2017). *gcl* is also required for germ cell transcriptional quiescence in flies (Leatherman et al., 2002). However, *gcl* is not required for germ cell specification in the mouse *Mus musculus* nor in the flour beetle *Tribolium castaneum*, but it is required for proper mouse spermatogenesis (Ansari et al., 2018; Kimura et al., 2003). Outside the germ line, *gcl* plays a role in establishing the anterior–posterior axis in *T. castaneum* (Ansari et al., 2018). Despite the wide divergence in their physiological roles, *gcl* homologs are highly conserved in amino acid sequence, with 45% similarity between homologs from *D. melanogaster* and *M. musculus*. In addition, these homologs appear to share at least some biochemical functions, since the mouse *gcl* homolog can partially restore germ cell formation in a *gcl*-null *D. melanogaster* embryo, though not to wild type numbers (Leatherman et al., 2000). *gcl* expression has also been observed in the embryonic germ line in other arthropod species (Ewen-Campen et al., 2013a; Gupta and Extavour, 2013).

However, to our knowledge, functional tests of whether *gcl* is required for insect germ line development or function remain limited to *D. melanogaster* and *T. castaneum*. Here, we examine the function of *germ cell-less* in the embryonic and adult germ lines of *O. fasciatus* and *G. bimaculatus*. Our data suggest that *gcl* is not required for germ cell specification nor for adult gametogenesis in either species. However, *gcl* mutant *O. fasciatus* embryos have significantly fewer germ cells than wild type. Knockdown of *Of*-*torso* does not affect germ cell number, suggesting that *gcl* in this species affects germ cell number through a different mechanism from that in *D. melanogaster*. Overall, these studies into *gcl* function across species inform our understanding of the evolution of gene function in germ line development.

## Materials and Methods

### Gene identification

#### germ cell-less

*Of*-*germ cell-less* is annotated as OFAS015300 on Scaffold 2113 of the current *O. fasciatus* genome assembly (Panfilio et al., 2019). *Gb*-*germ cell-less* is annotated as GBIM03474 on Scaffold 14 of the current *G. bimaculatus* genome assembly (Ylla et al., 2021). We amplified both genes from mixed stage embryonic cDNA of the respective species to verify the full-length coding sequences.

#### Of*-*torso

A tBLASTn search was performed on the *O. fasciatus* genome (Panfilio et al., 2019) using *D. melanogaster* Torso as a query, and a reciprocal BLASTp was performed on the top results with e-value < 1e–40. Of the initial results, OFAS005359 on Scaffold 336 returned *torso* orthologs from other insects. A reciprocal BLASTp search using the adjacent upstream annotated gene, OFAS005360, also returned *torso* orthologs from other insects. PCR was performed from mixed stage embryonic cDNA to confirm that OFAS005359 and OFAS005360 correspond to the same transcript. To obtain full-length sequence, RACE was performed using the SMARTer RACE 5’/3’ Kit (Takara Bio 634859) following manufacturer’s instructions.

The initial tBLASTn search using *D. melanogaster* Torso as a query also returned OFAS025175 as a result. A reciprocal BLASTp search using the predicted peptide sequence of this hit returned *fgf receptor* orthologs from other insects. Since the FGF receptor is a family of receptor tyrosine kinases closely related to Torso (Duncan et al., 2013), we wanted to include OFAS025175 for phylogenetic analysis. Therefore, we performed RACE using the SMARTer RACE 5’/3’ Kit to obtain the full-length sequence of this gene. Full length sequences for both *Of*-*torso* and *Of*-*fgfr* have been deposited at GenBank with accession numbers PV361543 and PV361544, respectively.

Phylogenetic analysis was performed using MEGA 11 (Tamura et al., 2021). Insect orthologs of Torso were aligned using MUSCLE (Edgar, 2004) along with orthologs of Ret oncogene and FGF receptor, which are the closest related families of receptor tyrosine kinases (Duncan et al., 2013). The multiple sequence alignment was used to generate a gene tree using the Maximum Likelihood method with the JTT substitution model (Jones et al., 1992), a discrete Gamma-distributed rate model and 1000 bootstrap replicates.

### Molecular biology

#### RNA extraction and cDNA synthesis

Embryos or tissues of interest were homogenized in TRIzol (Invitrogen 15596026), and RNA was extracted with phenol–chloroform extraction and isopropanol precipitation. cDNA was synthesized with SuperScript III First-Strand Synthesis SuperMix for qRT-PCR (Invitrogen 11752250) following manufacturer’s instructions.

#### Cloning and in vitro transcription

Fragments of genes of interest were PCR-amplified from embryonic cDNA and each cloned into the pGEM-T Easy vector (Promega A137A) using manufacturer’s instructions. A fragment of *gfp* was subcloned from an existing plasmid containing the *gfp* sequence. Insertion was confirmed by Sanger sequencing. Primers used are listed in Supplemental Table 1.

For *in situ* probe synthesis, gene fragments were PCR-amplified from plasmids using T7 and Sp6 primers. *In vitro* transcription was performed using T7 RNA polymerase (Promega P2075) or Sp6 RNA polymerase (Roche 10810274001) with DIG RNA labeling mix (Roche 11277073910). 100 ng of template was used for a 20-µl reaction run at 37°C overnight. RNA was purified with LiCl/ethanol precipitation, then resuspended in 20 µl DEPC-treated water. Concentration was measured using a NanoDrop Spectrophotometer (ThermoFisher), and the sample was diluted to 100 ng/µl with hybridization buffer (50% formamide, 1x phosphate-buffered saline (PBS), 5x saline–sodium citrate (SSC), 0.1% Triton X-100, 0.1% CHAPS, 5 mM EDTA, 50 µg/mL heparin, 1 µg/mL yeast tRNA). Probes were stored at −20°C.

For double-stranded RNA (dsRNA) synthesis, gene fragments were PCR-amplified from plasmids using a forward T7 primer and a reverse primer containing both T7 and Sp6 promoter sequences to generate a template for *in vitro* transcription with the T7 promoter sequence on both ends. *In vitro* transcription was performed using the MEGAscript T7 Transcription Kit (Invitrogen AM13345). 100 ng of template was used for an 80-µl reaction run at 37°C overnight. RNA was purified with LiCl/ethanol precipitation, then resuspended in 100 µl DEPC-treated water. To anneal the two strands, this solution was heated to 95°C on a heat block for 5 minutes, then allowed to cool slowly by leaving the tube on the heat block with the power switched off. A 1:10 dilution of annealed RNA was analyzed using a NanoDrop Spectrophotometer (ThermoFisher) to measure its concentration and purity, then run on a 1% agarose gel to confirm its size and integrity. For embryonic RNAi, dsRNA was diluted in DEPC-treated water to 2 µg/µl (*O. fasciatus*) or 5 µg/µl (*G. bimaculatus*), with 0.05% phenol red. For *G. bimaculatus* adult RNAi, dsRNA was diluted to 7.5 µg/µl in DEPC-treated water.

#### CRISPR/Cas9-mediated mutagenesis

Alt-R S.p. Cas9 Nuclease V3 (cat. #1081059) and Alt-R CRISPR-Cas9 tracrRNA (cat. #1072533) were ordered from IDT, along with gene-specific Alt-R CRISPR-Cas9 crRNA. tracrRNA and crRNA were resuspended to 100 µM in Nuclease-Free Duplex Buffer (IDT cat. #11-01-03-01). 5 µl of each RNA were combined in a PCR tube, heated to 95°C for 5 minutes, and allowed to cool to room temperature to form 50 µM gRNA duplex. 3 µl gRNA and 2 µl Cas9 protein (10 µg/µl or ∼62 µM) were combined in a PCR tube and incubated at room temperature for 10 minutes to form Cas9/gRNA ribonucleoprotein complex (RNP) at roughly 25 µM. Finally, RNP was diluted tenfold in 1x PBS with 0.05% phenol red.

### Of-Vasa antibody generation

A rabbit polyclonal antibody was raised against a recombinant *Of*-Vasa protein fragment (McGill Biology CIAN facility, Canada). The sequence encoding the 565 C-terminal amino acids of *Of*-Vasa, the stop codon and the first 96 base pairs of 3’ UTR (GenBank accession KC261571 (Ewen-Campen et al., 2013b)) was cloned into the pET151/dTOPO expression vector (Invitrogen K151-01), thus introducing an N-terminal 6x-polyhistidine tag. Protein expression was induced in *E. coli* BL21 (DE3) by addition of IPTG to a final concentration of 0.25 mM and shaking at 30°C for four hours. The overexpressed proteins were highly insoluble, so the cell pellets were disrupted by sonication in non-denaturing buffer, and the soluble fraction was separated by centrifugation and discarded. The pellet was resuspended in 8 M urea (crude prep). Crude preps were further purified by SDS gel-purification. Protein bands were excised from the gel after reverse staining with zinc chloride and the proteins were collected by electroelution. Acetone precipitation was performed, and the precipitated protein was washed and resuspended in 8 M urea. Protein concentration was adjusted to 1–2 mg/mL for injection. Rabbits were injected using a standard 80-day protocol with three boosts after the initial injection; the final bleed was performed after 87 days. The serum was processed by addition of sodium azide to 0.02% (w/v). The antibody was affinity-purified against bacterially expressed *Of*-Vasa protein.

### Embryo microinjection

Needles for injection were made by pulling glass capillaries (World Precision Instruments 1B100-4) with a Sutter P-97 Flaming/Brown micropipette puller. Needle tips were manually opened with fine forceps just prior to injection. Eggs were injected with a Narishige IM300 microinjector.

*O. fasciatus* were allowed to lay eggs in dry cotton for four hours, after which eggs were picked out of the cotton for injection. Two glass microscope slides were taped together with double-sided tape to create a ledge with exposed double-sided tape against which embryos were lined up. Eggs were then submerged in tap water for at least 20 minutes to soften the eggshell before injection. Slides with injected eggs were kept at 28°C to continue development until the desired age.

*G. bimaculatus* were allowed to lay eggs in wet cotton for four hours, after which eggs were washed out of the cotton for injection. Eggs were aligned in grooves (570 µm wide × 2930 µm long × 650 µm deep) made in 2% agarose in Milli-Q water using a custom mold as previously described (Donoughe et al., 2018). Eggs were injected while submerged in 1x PBS. After injection, the solution was replaced with fresh 1x PBS. Injected eggs were kept submerged in 1x PBS at 28°C overnight. The following day, dead embryos, if any, were removed, and the remaining eggs were transferred onto damp Kimwipes in a covered Petri dish kept at 28°C to continue development. Three days after injection, eggs were submerged in tap water again and monitored every day until they reached the desired stage for dissection.

### Embryo dissection and fixation

*O. fasciatus* eggs were stuck to the bottom of a 60-mm Petri dish with double-sided tape and a slit was created in the eggshell with a microscalpel. 1x PBS with 4% paraformaldehyde (PFA) was then poured into the dish to fix the embryos for 30 minutes at room temperature, followed by at least three washes with PBT (1x PBS with 0.1% Triton X-100). Embryos were then dissected free of their yolk in PBT with fine forceps. Embryos were washed at least three times with PBT at room temperature before continuing with staining or dehydration in methanol.

*G. bimaculatus* embryos were dissected free of their yolk in 1x PBS with fine forceps and fixed overnight in 1x PBS with 4% PFA at 4°C. The following morning, they were washed at least three times with PBT at room temperature before continuing with staining or dehydration in methanol.

### G. bimaculatus adult RNA interference (RNAi)

2 µl dsRNA was injected into the abdomen of an adult cricket with a Hamilton syringe through the base of a metathoracic leg. Gonads were dissected seven days after injection for analysis.

### Colorimetric in situ hybridization

Dissected embryos were stored in methanol at −20°C overnight or longer before *in situ* hybridization. After rehydration in PBT and washing in hybridization buffer, embryos were hybridized overnight in a 70°C water bath with 1 ng/µl probe in hybridization buffer. The following day, the following series of 20-minute washes were performed at 70°C: once in hybridization buffer, three times in wash 1 (2x SSC, 0.1% CHAPS) and three times in wash 2 (0.2x SSC, 0.1% CHAPS). Embryos were then washed three times for ten minutes each at room temperature in KTBT (1x Tris-buffered saline with KCl and 0.1% Triton X-100) followed by an hour-long incubation at room temperature in *in situ* blocking solution (1x PBS, 0.1% Tween-20, 5% NGS, 2 mg/mL bovine serum albumin, 1% dimethyl sulfoxide). Embryos were incubated for two hours at room temperature with sheep anti-digoxigenin–AP (Roche 11093274910), diluted 1:2000 in *in situ* blocking solution, then washed at least four times over two hours with KTBT and incubated in KTBT overnight at 4°C. The following day, embryos were washed with NTMT (100 mM Tris pH 9.5, 100 mM NaCl, 50 mM MgCl_2_, 0.1% Tween-20), then developed in the dark at 37°C with NBT/BCIP in NTMT with 5% polyvinyl alcohol. Development was monitored at least every hour under a dissection microscope. To stop development of the enzymatic reaction, embryos were washed in PBT, incubated with 5 ng/µl Hoechst 33342 in PBT at room temperature for an hour, post-fixed in 4% PFA in PBS, then incubated in 70% glycerol before mounting in 100% glycerol.

### Immunohistochemistry

Embryos were incubated in blocking solution (PBT with 5% normal goat serum (Jackson ImmunoResearch 005-000-121)) for one hour at room temperature, then incubated overnight at 4°C with primary antibodies diluted in blocking solution. The following day, embryos were washed for at least four times over four hours with PBT, incubated for an hour in blocking solution, then incubated overnight at 4°C with secondary antibodies diluted in blocking solution. Finally, embryos were washed in PBT, post-fixed in 4% PFA in PBS, then incubated in 70% glycerol before mounting in glycerol. Antibody staining was also performed with a no-primary control.

Primary antibodies used were: rabbit anti-*Gb*-Piwi (Ewen-Campen et al., 2013a) at a final concentration of 1:300; rabbit anti-*Of*-Vasa at 1:100; mouse anti-α-tubulin (DM1A; Sigma T9026) at 1:100; rabbit anti-β-catenin (Sigma C2206) at 1:500; and mouse anti-RNA polymerase II (H5; Covance MMS-129R) (Bregman et al., 1995; Warren et al., 1992) at 1:100. Secondary antibodies used were goat anti-rabbit conjugated to Alexa Fluor 647 (Invitrogen A-21246), goat anti-rabbit Alexa Fluor 488 (Invitrogen A-11008), goat anti-rabbit Alexa Fluor 568 (Invitrogen A-11011), donkey anti-mouse Alexa Fluor 488 (Invitrogen A-21202), and goat anti-mouse Alexa Fluor 647 (Invitrogen A-21236). All secondary antibodies were used at a final concentration of 1:400 along with Hoechst 33342 (Sigma B2261) at 5 ng/µl.

Embryos were imaged on a Nikon CSU-W1 spinning disk inverted confocal microscope with 10x air or 20x air objectives or a Zeiss AxioObserver inverted compound microscope with 10x air or 20x air objectives. Images were analyzed in Fiji.

### Quantitative RT-PCR

Quantitative RT-PCR was performed on an AriaMx Real-time PCR system (Agilent) using PerfeCTa SYBR Green SuperMix, Low ROX (Quantabio 95056). Fold-change differences were calculated using the ΔΔC_t_ method for RNAi experiments. Expression levels were normalized against *Of*-*gapdh* or *Gb*-*β-tubulin*. All primers used are listed in Supplemental Table 1.

## Results

### germ cell-less expression across insect species

During oogenesis in *D. melanogaster*, Gcl protein is detectable at the nuclear membrane of nurse cells and oocytes (Moore et al., 2009). In addition, *gcl* transcripts are transcribed by nurse cells, transported into the oocyte and deposited at the oocyte posterior, where they remain as part of the germ plasm into early embryogenesis (Jongens et al., 1992). These transcripts are translated into protein starting at the cleavage stages of embryogenesis (Jongens et al., 1992). Gcl protein is localized to the nuclear membranes of germ cells after they form and remains detectable in germ cells until mid-gastrulation, when germ cells enter the primordial midgut (Jongens et al., 1992). We asked whether *gcl* is also expressed in other insect embryos at the time and place of germ cell specification.

We (Ewen-Campen 2013) and others (Butt 1949) have qualitatively described the first appearance of primordial germ cells in blastoderm stage *O. fasciatus* embryos, and their proliferation and migration during embryogenesis. However, to our knowledge there has been no embryonic developmental staging system established for this insect. We therefore we defined stages based on major embryological, morphogenetic and morphological milestones, for describing embryonic development from egg-laying through to katatrepsis, with special emphasis on the development of germ cells (Supplemental Figure 1; Supplemental Table 2). We assessed *gcl* transcript levels using quantitative RT-PCR on *O. fasciatus* embryos collected in 12-hour windows spanning the first four days of embryogenesis. *Of*-*gcl* transcripts were detected at the highest levels 24–36 hours after egg-lay at 28°C (Figure 1A). This window corresponds to the late blastoderm stage (stage 3), when germ cells are first distinguishable (Ewen-Campen et al., 2013b), and the beginning of germ band elongation (stages 4–5). We speculate that this peak in expression is likely due to zygotic rather than maternal expression, since it is higher than expression in freshly laid eggs. However, we were unable to detect *Of*-*gcl* transcripts by *in situ* hybridization above background levels in embryos at any stage of development, nor in adult ovaries or testes (Supplemental Figure 2A–B).

**Figure 1.**
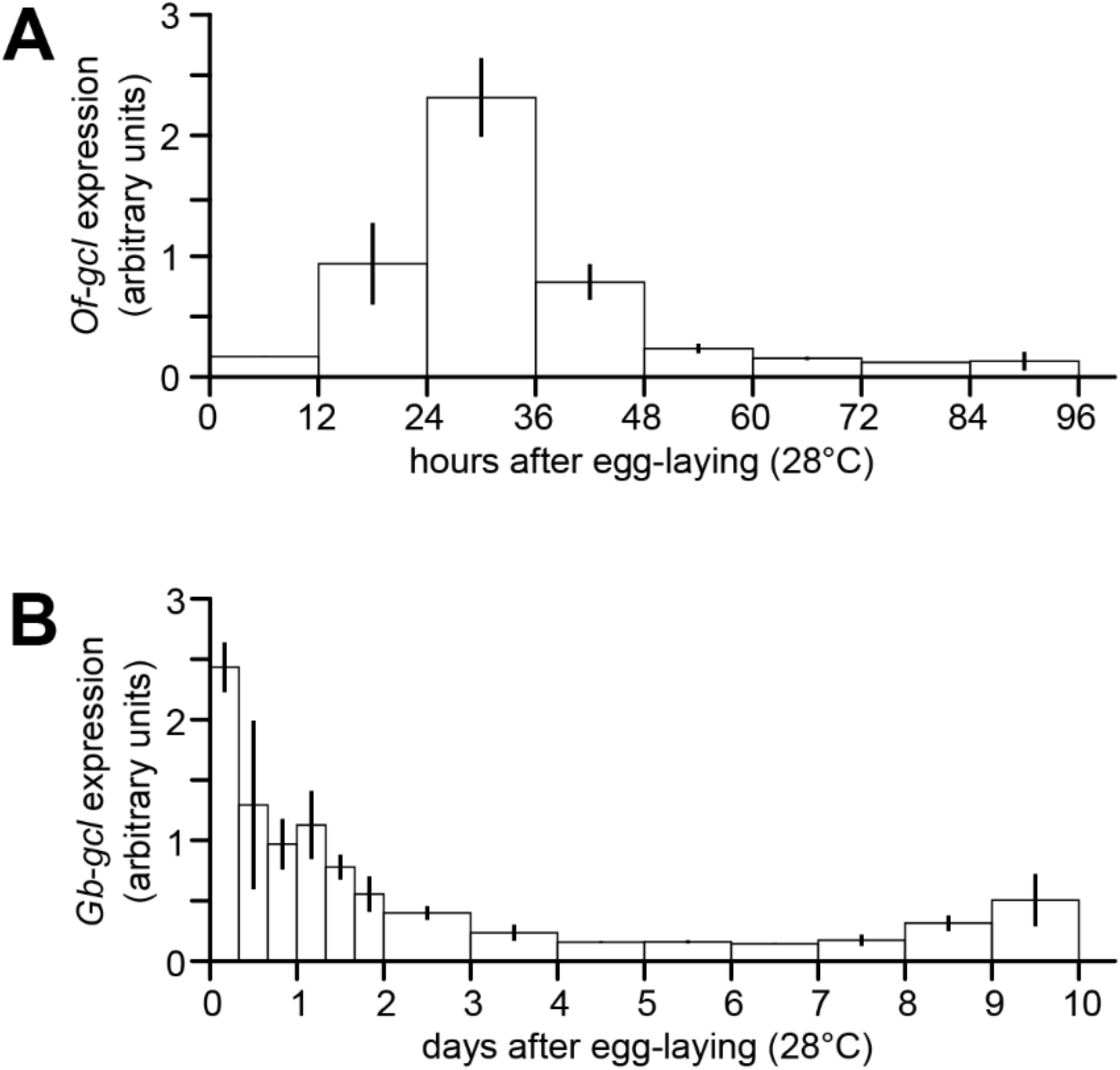
*gcl* expression is detectable by qPCR in *O. fasciatus* (A) and *G. bimaculatus* (B) embryos. C_q_ values were normalized against *Of*-*gapdh* or *Gb*-*tubulin* expression and averaged over four biological replicates for each time point. Error bars represent standard deviations. The width of each bar represents the length of time for each embryo collection.

We similarly assessed *gcl* transcript levels using qRT-PCR on *G. bimaculatus* embryos collected in 8-hour windows spanning the first two days of embryogenesis and 24-hour windows spanning the following eight days of embryogenesis. *Gb*-*gcl* transcripts were detectable at the highest levels in the first eight hours after egg-lay (Figure 1B), suggesting maternal deposition of *Gb*-*gcl* transcript. Consistent with this hypothesis, *Gb*-*gcl* transcript was detectable by *in situ* hybridization in ovaries but not during embryogenesis (Supplemental Figure 2C–D).

Although we were unable to detect localized *gcl* expression in either *O. fasciatus* or *G. bimaculatus* embryos, we still sought to determine whether the gene’s documented roles in germ cell specification (*D. melanogaster*) or adult spermatogenesis (*M. musculus*) might be conserved (Jongens et al., 1992; Kimura et al., 2003). To this end, we analyzed *gcl* loss of function conditions in embryos and adult gonads of both species. *gcl* is also required for transcriptional quiescence in germ cells in *D. melanogaster* (Leatherman et al., 2002). However, we could not detect any reduction in transcriptional activity in germ cells relative to somatic cells of either *O. fasciatus* or *G. bimaculatus* (Supplemental Figure 3), so we did not assess this phenotype in our functional studies of *gcl*.

### Of-gcl mutation decreases germ cell number in milkweed bug embryos

To determine whether *gcl* is required for germ cell specification in *O. fasciatus*, we generated an *Of*-*gcl* putative protein-null allele with CRISPR/Cas9-mediated mutagenesis (Figure 2A–B). This allele contains a 2-bp insertion at amino acid residue 14. The resulting frameshift leads to a premature stop codon, resulting in a predicted peptide of 30 amino acid residues, compared with the 478-residue wild type protein. Surprisingly, individuals homozygous for this mutant allele were viable and fertile, and we have maintained a stable mutant line for at least twelve generations at the time of writing. There were no detectable morphological defects in adult ovaries or testes from this mutant line, nor any evidence of abnormal gametogenesis in either males or females (Supplemental Figures 4–5).

**Figure 2.**
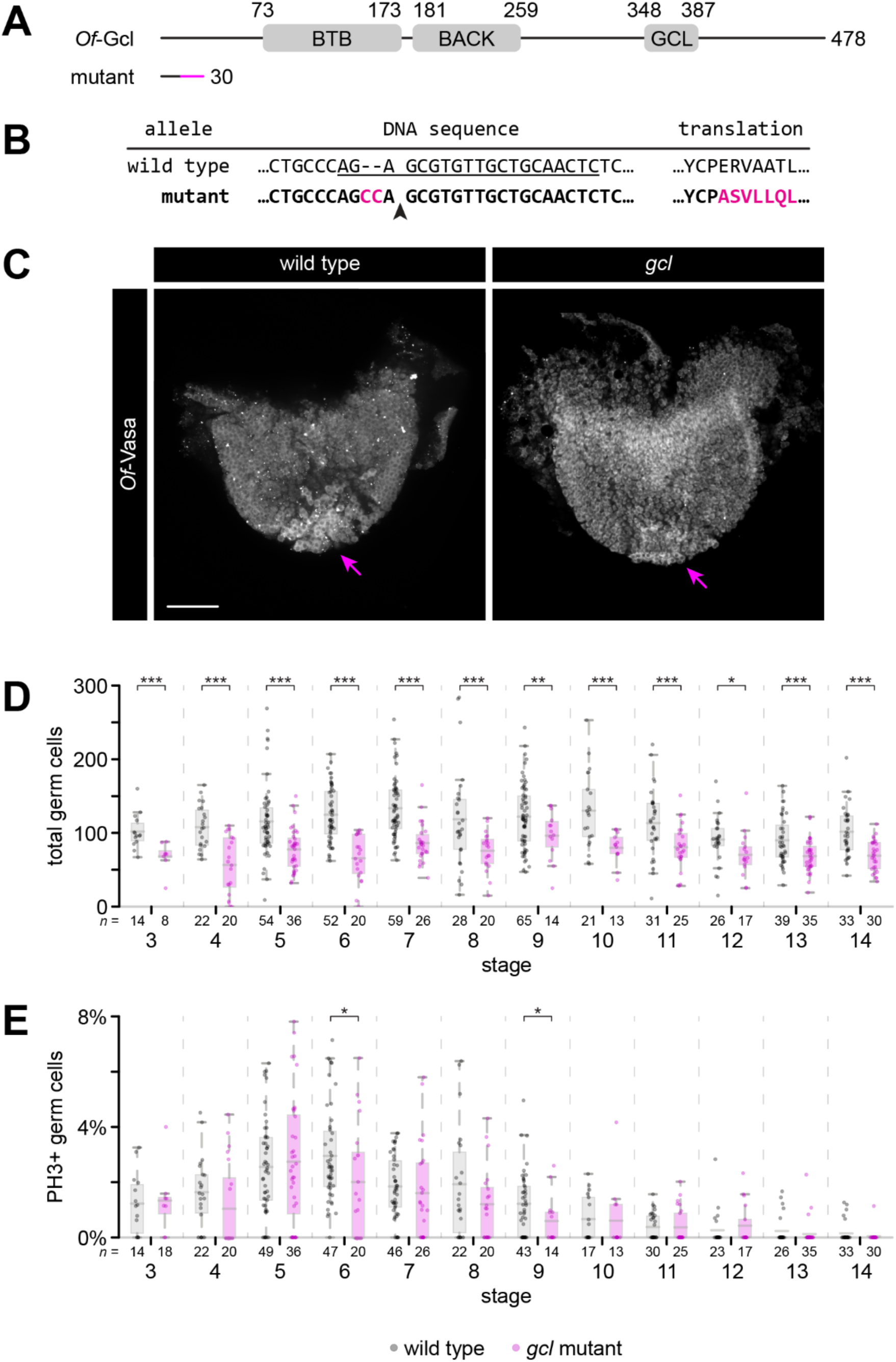
*Of*-*gcl* mutation decreases germ cell number in embryos. (**A**) Predicted domain architecture of *Of*-Gcl and predicted mutant protein. Wild type *Of*-Gcl has two predicted folded domains (BTB and BACK domains) as well as a conserved motif called the GCL domain. The predicted mutant protein contains none of these domains and is truncated after residue 30. (**B**) *Of*-*gcl* mutant allele contains a two base pair insertion, creating a frameshift and leading to a predicted truncated protein. Guide sequence is underlined; arrowhead marks predicted Cas9 cut site. Inserted bases and predicted mutated residues are indicated in magenta. (**C**) Representative antibody stains against *Of*-Vasa to mark germ cells in stage 3 wild type and *gcl* mutant embryos. Mutant embryos still have germ cells at the expected location, indicated by arrows. Scale bar = 100 µm. (**D**) Germ cell counts across embryogenesis in wild type and *gcl* mutant embryos. Mutant embryos have significantly fewer germ cells per embryo compared to wild type. (**E**) Proportion of PH3^+^ germ cells across embryogenesis in wild type and *gcl* mutant embryos. There is no significant difference in the proportion of proliferating germ cells between wild type and *gcl* mutant embryos. All statistical tests are non-parametric bootstrap comparisons. Embryonic stages are defined in Supplemental Figure 1 and Supplemental Table 3. * *p* < 0.05; ** *p* < 0.01; *** *p* < 0.001

In *O. fasciatus*, germ cells are first detectable in stage 3 on the interior of the cellular blastoderm, forming part of an invaginating structure called the posterior pit (Ewen-Campen et al., 2013b). The germ cells remain at the posterior end of the germ band as it forms by invaginating into the egg (Ewen-Campen et al., 2013b) (Figure 2B; Supplemental Figure 1). At stage 3, *gcl* mutant embryos had on average 35% fewer germ cells than wild type embryos (*p* < 0.001; *n* = 14 wild type, 8 *gcl* embryos; Figure 2C–D; Supplemental Table 3). Germ cell number remained significantly lower in *gcl* mutant embryos relative to wild type embryos throughout all embryonic stages examined (Figure 2D). We detected a marginal decrease in germ cell proliferation in *gcl* mutant embryos relative to wild type only at stages 6 and 9, as measured by the proportion of germ cells with detectable phospho-histone 3 (Figure 2E; Supplemental Table 3). However, there was no detectable difference in germ cell proliferation between the two genotypes at any other embryonic stage examined (Figure 2E; Supplemental Table 3). We were also unable to detect cleaved caspase 3-dependent cell death in stages 3–4 of either genotype (*n* = 27 wild type, 23 *gcl* embryos).

It is formally possible that *gcl* loss of function affects the expression of Vasa protein, which we use as an experimental proxy for identifying germ cells. However, we noted that the Vasa-positive cells in *gcl* mutant embryos expressed Vasa at comparable levels to wild type embryos. Thus, while we cannot rule out the possibility that germ cells that do not express detectable Vasa may be present in our *gcl* loss of function embryos, we believe it unlikely that lack of *gcl* would eliminate Vasa expression in some germ cells without significantly affecting expression in others. Therefore, we favor the interpretation that the reduction in Vasa-positive cells in *gcl* mutant embryos reflects a reduction in germ cell number. Given that the discrepancy in germ cell number did not appear to be the result of differences in cell proliferation or cell death, these data suggest that *Of*-*gcl* is involved in, but not strictly necessary for, one or more of germ cell specification, formation or maintenance.

In *D. melanogaster*, Gcl promotes germ cell fate by targeting the receptor Torso for degradation, and *torso* RNAi in *gcl* mutant flies restores germ cell numbers to wild type levels (Pae et al., 2017). To determine whether this mechanism of germ cell number determination is conserved in *O. fasciatus*, we asked whether *torso* loss of function also affects germ cell number in this species. Despite previous reports that the *O. fasciatus* genome lacks a *torso* ortholog (Skelly et al., 2019; Weisbrod et al., 2013), we identified a high-confidence homolog in the published genome assembly (Panfilio et al., 2019) (Figure 3A). Knockdown of *Of*-*torso* by embryonic RNAi did not affect germ cell number in stages 3–4 in either wild type or *gcl* mutant backgrounds (Figure 3B–C). qPCR analysis of injected embryos confirmed that RNAi was effective at reducing *torso* transcript levels to 50% relative to controls (Supplemental Figure 6A). This result suggests that the mechanism through which *gcl* controls germ cell number is not conserved between *O. fasciatus* and *D. melanogaster*.

**Figure 3.**
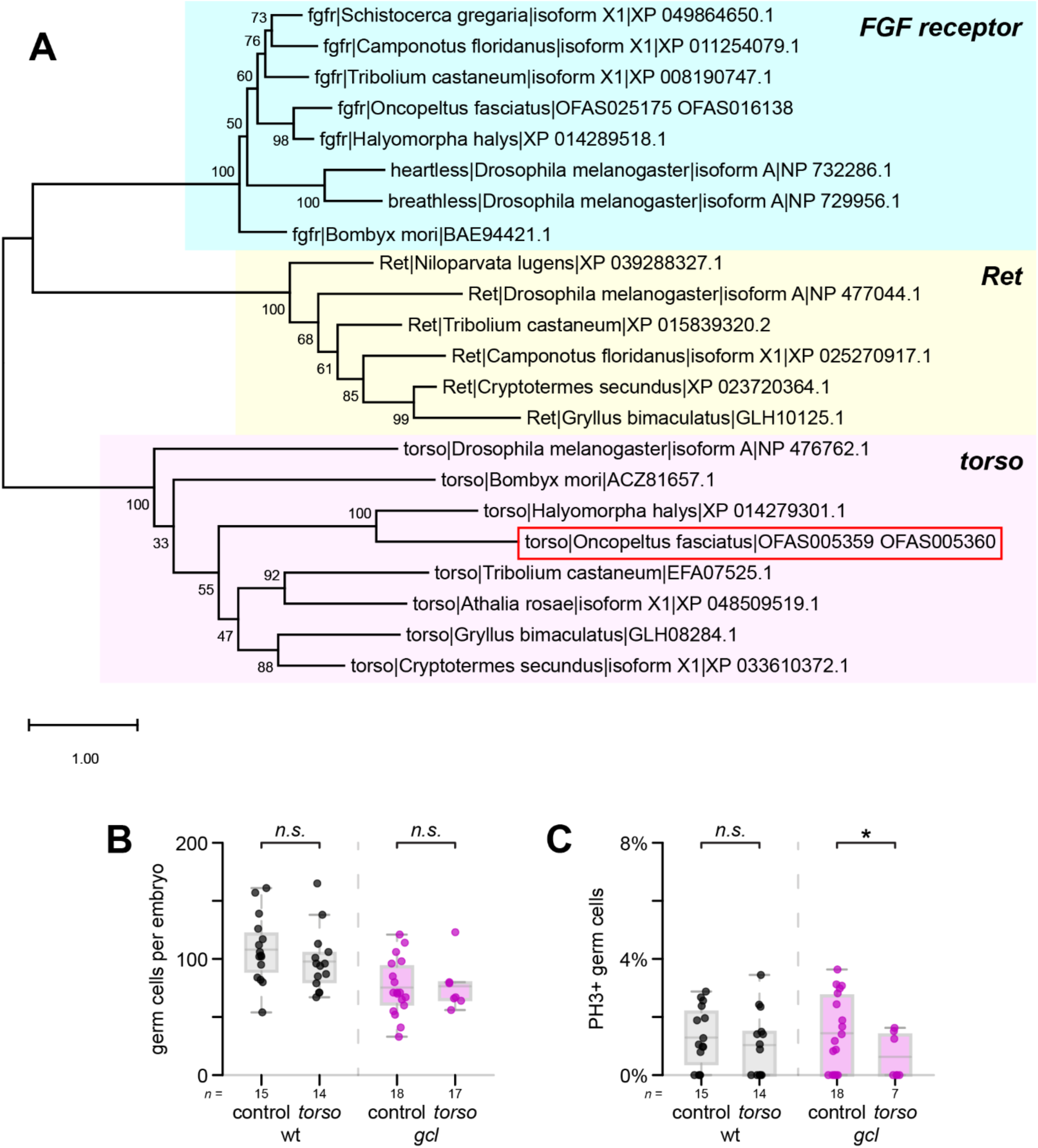
*Of*-*torso* knockdown does not affect embryonic germ cell number. (**A**) Phylogenetic analysis of *torso* and related receptor tyrosine kinases (*Ret oncogene* and *FGF receptor*) across insects. Each clade is individually highlighted for clarity. The putative *torso* homolog from *O. fasciatus* (OFAS005359/OFAS0005360) falls within the *torso* clade. (**B**) Germ cell counts in stage 3–4 *Of*-*torso* RNAi embryos, performed in wild type and *gcl* mutant backgrounds. *torso* knockdown does not affect germ cell number in either background. (**C**) Proportion of PH3^+^ germ cells in stage 3–4 *Of*-*torso* RNAi embryos, performed in wild type and *gcl* mutant backgrounds. *torso* knockdown marginally reduces germ cell proliferation in *gcl* mutant embryos but not in wild type embryos. All statistical tests are non-parametric bootstrap comparisons. * *p* < 0.05; ** *p* < 0.01; *** *p* < 0.001

### Gb-gcl knockdown does not affect germ cell number in cricket embryos

To determine whether *gcl* is required for germ cell specification in *G. bimaculatus*, we knocked down *gcl* expression using embryonic RNAi. We found no significant difference in total germ cell number in stage 8 embryos under *gcl* knockdown relative to *gfp* controls, nor did we detect any significant difference in germ cell number in any segment (Figure 4). qPCR analysis of injected embryos confirmed that RNAi was effective at reducing *gcl* transcript levels to 34% relative to controls (Supplemental Figure 6B). It is possible that knockdown efficiency is not high enough with RNAi to affect germ cell specification. However, we note that we have previously detected significant changes in germ cell number with comparable or lower knockdown efficiencies (Barnett et al., 2019; Donoughe et al., 2014; Nakamura and Extavour, 2016). Therefore, we speculate that the role of *Gb*-*gcl* in germ cell specification or maintenance, if any, must be subtle.

**Figure 4.**
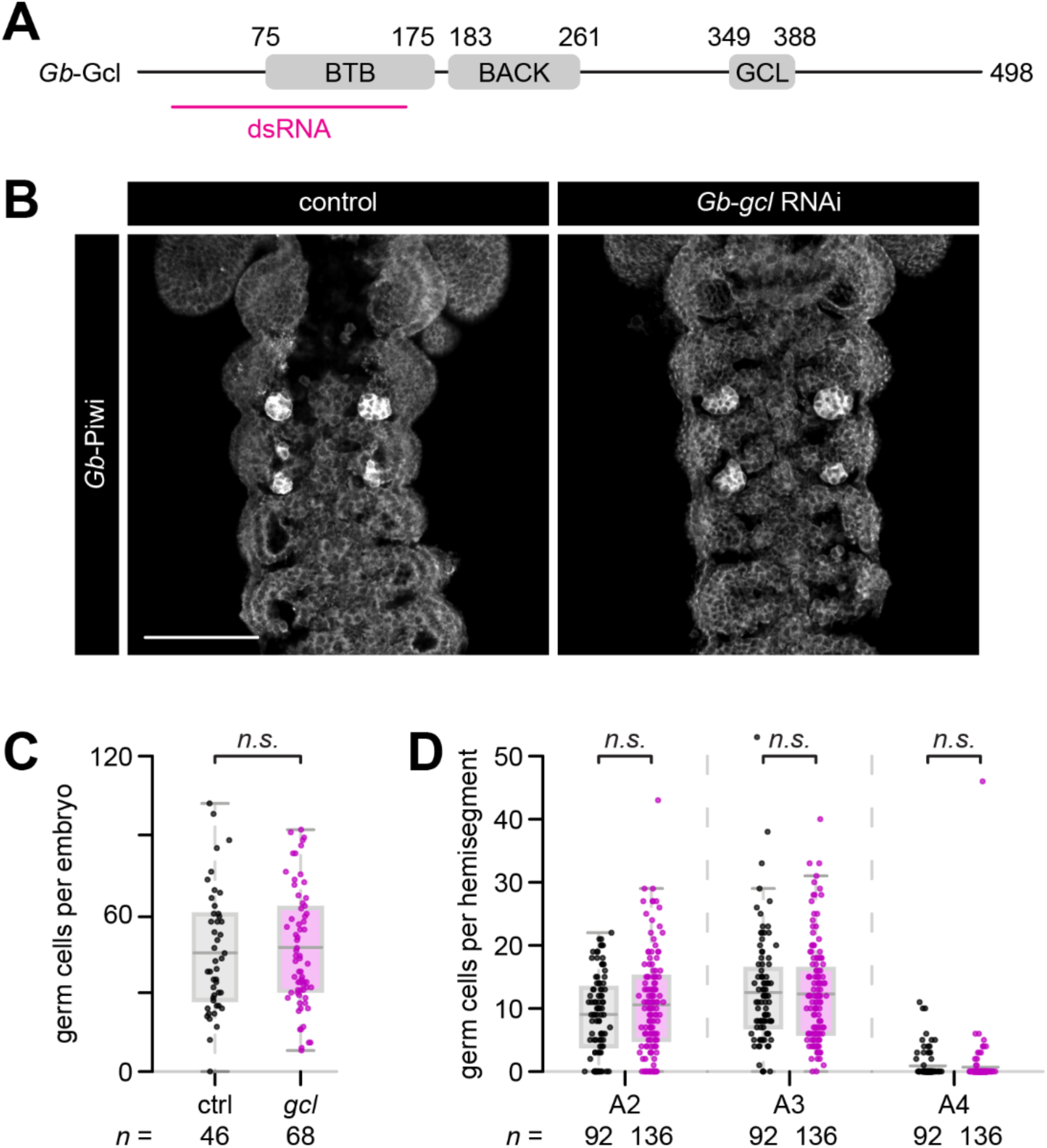
*Gb*-*gcl* knockdown does not affect embryonic germ cell number. (**A**) Predicted domain architecture of *Gb*-Gcl. *Gb*-Gcl has two predicted folded domains (BTB and BACK domains) and a conserved motif called the GCL domain. The fragment region used for dsRNA synthesis is marked. (**B**) Representative antibody stains against *Gb*-Piwi to mark germ cells in stage 8 control and *gcl* knockdown embryos. Embryos in which *Gb*-*gcl* has been knocked down still have germ cells in the expected positions. Scale bar = 100 µm. (**C**) Germ cell counts in stage 8 *Gb*-*gcl* RNAi embryos. Germ cell numbers are not significantly different in *Gb*-*gcl* knockdown embryos relative to controls. Each point represents one embryo. (**D**) Germ cell counts in stage 8 *Gb*-*gcl* RNAi embryos, divided into hemisegments (*i.e.*, the left or right half of one segment). *Gb*-*gcl* knockdown does not affect the number of germ cells in any individual segment. Each point represents a hemisegment. All statistical tests are non-parametric bootstrap comparisons.

Next, we asked whether *gcl* has any role in adult gametogenesis in *G. bimaculatus* by performing RNAi in adult crickets. We could not detect any morphological or gametogenesis defects in either ovaries or testes (Supplemental Figures 4-5). These data suggest that *gcl* is not required for adult gametogenesis in this cricket.

## Discussion

Our analysis of *gcl* loss of function in embryos of two different hemimetabolous insects suggests that *gcl* is not required for germ cell specification in these species, in contrast to the gene’s importance in germ cell formation in *D. melanogaster*. However, *Of*-*gcl* knockout embryos have significantly fewer germ cells than wild type embryos, suggesting that the gene may be involved in germ cell specification or maintenance in this bug, even if it is not strictly required.

Gcl protein suppresses the somatic cell fate in *D. melanogaster* by targeting the receptor tyrosine kinase Torso for degradation (Colonnetta et al., 2021; Pae et al., 2017). Germ cell number can thus be modulated in flies by up- or downregulating Torso activity (Pae et al., 2017). However, RNAi knockdown of *torso* in *O. fasciatus* does not affect germ cell number in either wild type or *gcl* mutant embryos (Figure 3). Therefore, we hypothesize that germ cell specification in this species does not require downregulation of Torso by Gcl, and that *Of*-*gcl* regulates germ cell specification through some other unknown pathway. Torso’s role in germ cell specification may have been co-opted from its ancestral functions in neuroendocrine signaling, just as it is thought to have been co-opted into its role in terminal patterning in flies (Duncan et al., 2013; Schoppmeier and Schröder, 2005).

In *D. melanogaster*, the interaction between Gcl and Torso proteins depends on sequence motifs present on both proteins (Pae et al., 2017). In Gcl protein, this motif is the GCL domain, which is conserved across insect homologs (Supplemental Figure 7) (Pae et al., 2017). Torso interacts with Gcl through a motif in the linker domain between its split kinase domains. This linker region is significantly longer in *Dm*-Torso compared with other insect homologs (Supplemental Figure 8), and the specific motif identified by Pae and colleagues (Pae et al., 2017) is not present in other homologs. The conservation of the GCL domain despite the lack of conservation in the Torso interaction motif may suggest that Gcl has some other unknown interaction partner, or that the Gcl–Torso interaction may not be as sequence-specific as originally thought. AlphaFold3 (Abramson et al., 2024) predicts potential interaction between Gcl and Torso homologs in both *O. fasciatus* and *G. bimaculatus* through the split kinase linker domain of Torso (Supplemental Figure 9). Further experiments will be needed to test whether these proteins interact *in vivo*, and to identify additional conserved interaction partners of Gcl.

In mice, *gcl* knockout results in abnormal nuclear morphology in the liver, pancreas and testis, though the only functional defect identified was impaired spermatogenesis (Kimura et al., 2003). Similarly, *gcl* is expressed at low levels in somatic tissues in flies including the gut, muscle and nervous system, but no defects have been reported in those tissues in *gcl*-null flies (Jongens et al., 1992; Robertson et al., 1999). We were also unable to detect any defects in spermatogenesis under either *Of*-*gcl* knockout or *Gb*-*gcl* knockdown conditions (Supplementa1 Figure 4). Given that the mouse *gcl* homolog can rescue germ cell formation in *gcl*-null flies (Leatherman et al., 2000), its biochemical functions are likely to be conserved across these species. We speculate that *gcl* has a ubiquitous but subtle cell biological role in maintaining nuclear morphology that only yields a phenotype in certain tissue contexts, such as in mouse spermatogenesis. Under this hypothesis, *gcl* then gained an additional role in germ cell specification in certain lineages of insects. In addition, the mechanism underlying this germ cell specification has diverged between insect lineages, perhaps through the evolution of new interaction partners such as Torso. This model reflects the complexity and variation present in the mechanisms of germ cell specification and their evolution, including the co-option of new genes with the shifting mechanisms of this process across animals.

## Author contributions

JAK: Conceptualization, Methodology, Formal Analysis, Investigation, Writing – Original Draft, Writing – Review & Editing, Visualization; BE-C: Resources; CGE: Conceptualization, Writing – Review & Editing, Supervision, Funding Acquisition

## Acknowledgments

The authors thank members of the Extavour lab for valuable discussions. This research was supported by National Science Foundation (NSF) Awards IOS-2220747 and IOS-0817678 to CGE, and an NSF Graduate Research Fellowship to BE-C. CGE is a Howard Hughes Medical Institute Investigator.

## Notes

### Competing Interest Statement

The authors have declared no competing interest.

### Summary of Updates

1. addition of new supplementary data providing quantitative evidence for efficacy of RNAi. 2. expanded discussion of interpretation of transcriptional activity data. 3. introduction of staging information earlier in the text for improved clarity.

