## Supplementary Material for "Divergence of *germ cell-less* roles in germ line development across insect species"

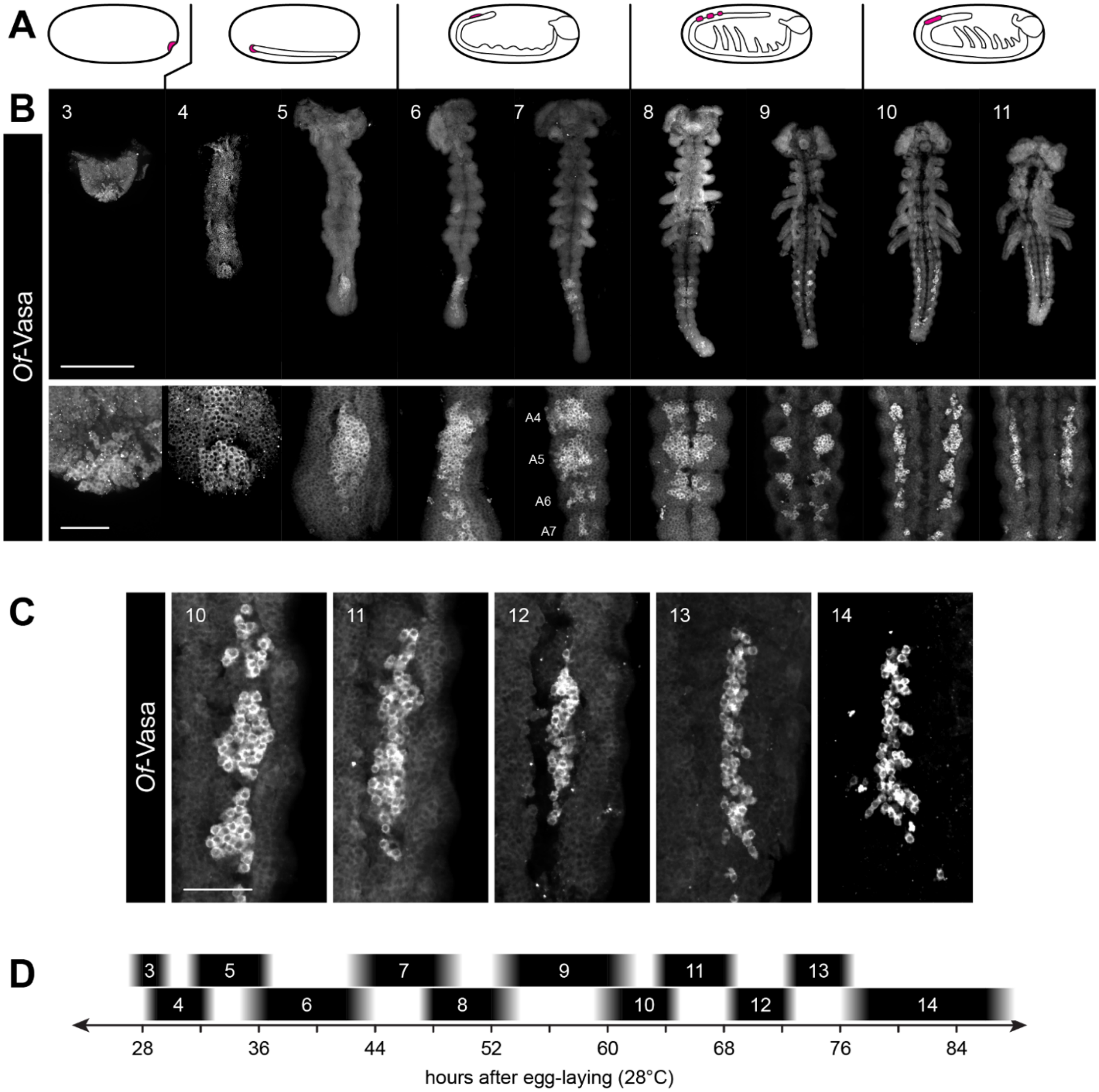


**Supplemental Figure 1. Wild type germ line development in *O. fasciatus*.**

(**A**) Schematics showing the position of the *O. fasciatus* embryo within the egg, roughly corresponding to the stages shown below in (**B**). Germ cells are indicated in magenta.

(**B**) Representative images of *Of*-Vasa antibody stains in stage 3–11 *O. fasciatus* embryos. Top row shows whole mount embryos, while the bottom row shows a higher magnification view of the germ cells. Scale bar = 500 µm in top row, 100 µm in bottom row.

(**C**) *Of*-Vasa antibody stains in stages 10–14 showing coalescence of germ cells into the embryonic gonad. Right gonad shown in all images. Scale bar = 50 µm.

(**D**) Timeline of embryonic stages for wild type development at 28°C, 60% humidity.

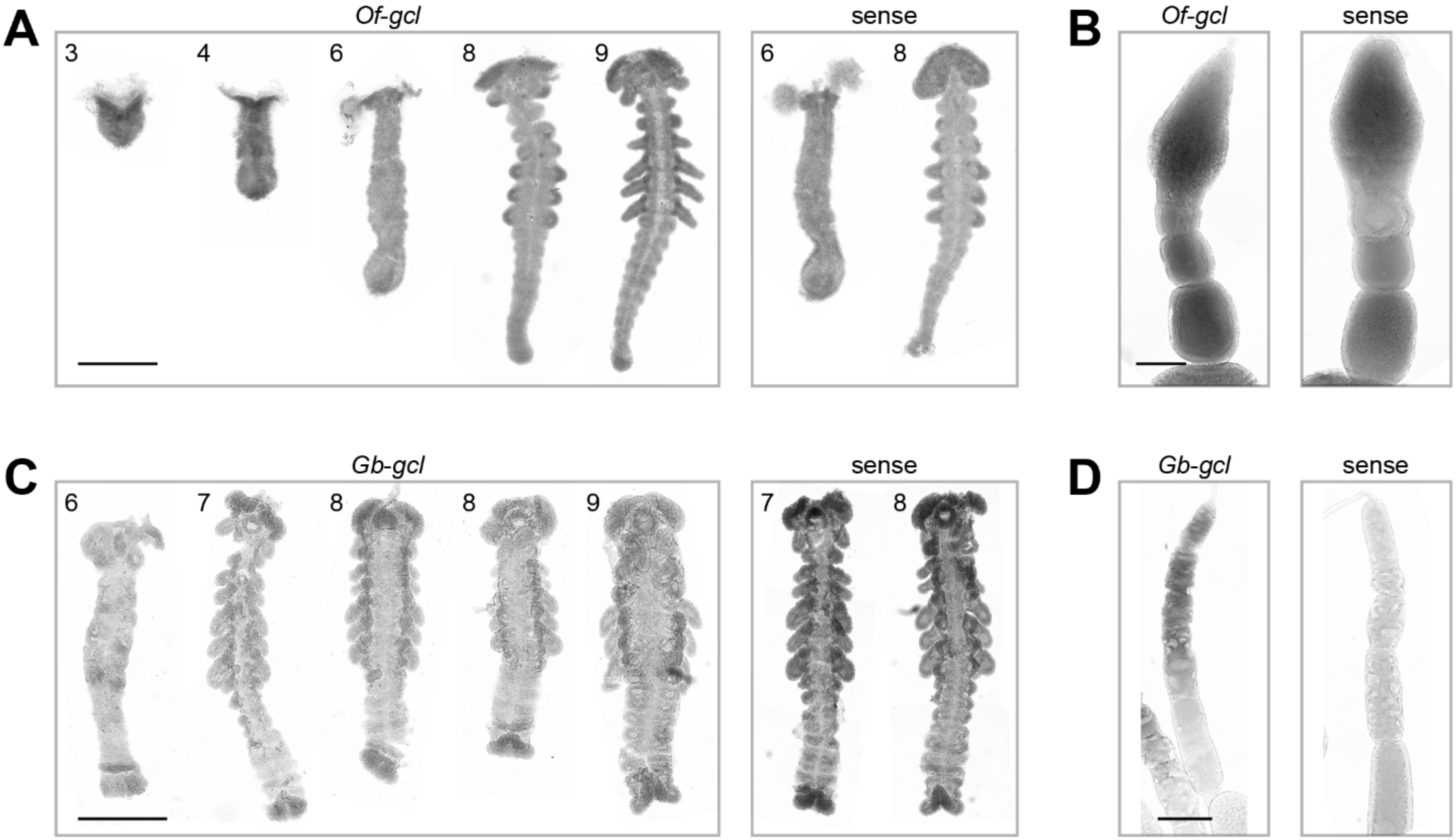


**Supplemental Figure 2. *gcl* is not detectable with *in situ* hybridization during embryogenesis in *O. fasciatus* or *G. bimaculatus*.**

(**A**–**B**) *In situ* hybridization against *Of-gcl* in embryos (C) and ovaries (D). Numbers indicate embryonic stages. Scale bars = 500 µm (embryos) or 200 µm (ovaries).

(**C**–**D**) *In situ* hybridization against *Gb-gcl* in embryos (E) and ovaries (F). Numbers indicate embryonic stages. Scale bars = 500 µm (embryos) or 200 µm (ovaries).

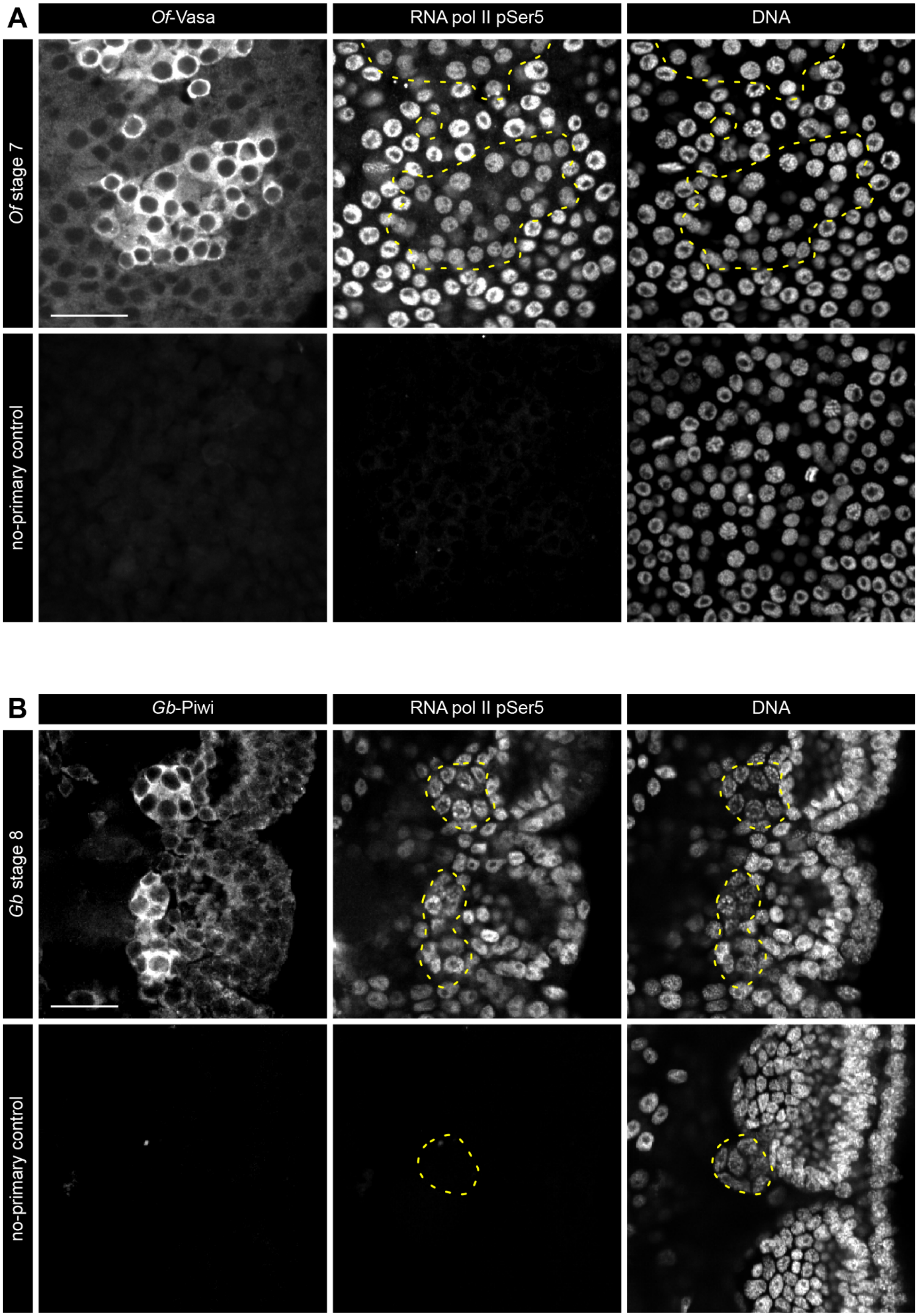
A
Of-Vasa
RNA pol II pSer5
DNA
Of stage 7
no-primary control
B
Gb-Piwi
RNA pol II pSer5
DNA
Gb stage 8
no-primary control

**Supplemental Figure 3** (previous page)**. Germ cells in *O. fasciatus* and *G. bimaculatus* are not transcriptionally quiescent.**

(**A**) Antibody stains in stage 7 *O. fasciatus* embryos against *Of*-Vasa to visualize germ cells and against hyperphosphorylated RNA polymerase II (H5 antibody) to visualize transcriptional activity. Embryos were counterstained with Hoechst 33342 to visualize nuclei. Scale bar = 25 µm.

(**B**) Antibody stains in stage 8 *G. bimaculatus* embryos against *Gb*-Piwi to visualize germ cells and against hyperphosphorylated RNA polymerase II (H5 antibody) to visualize transcriptional activity. Embryos were counterstained with Hoechst 33342 to visualize nuclei. Scale bar = 25 µm.

In both species, hyperphosphorylated RNA polymerase II is detectable in germ cells (indicated by dashed yellow lines) at levels higher than the background seen in the no-primary control, suggesting transcriptional activity in germ cells. While the fluorescence intensity of the H5 antibody signal may appear reduced in some germ cell nuclei relative to somatic nuclei in both species, we note that all images presented are single 1 µm optical sections, and that signal quantification through the entire 3D volume of each nucleus would be needed to determine whether there is indeed a significant difference in signal among germ cells, among somatic cells, or between germ cells and somatic cells. Apparent differences in fluorescence intensity may be related to the distinct chromatin architecture in germ cell nuclei relative to somatic nuclei, rather than by a significant reduction in transcriptional activity. Orthogonal methods such as EU labeling will be required in future studies to determine whether transcriptional activity is reduced in germ cells relative to somatic cells.

Even if that were the case, our data stand in contrast to *D. melanogaster*, in which no H5 signal above background levels has been reported in primordial germ cells from their formation in stage 5 until stage 7, when they are internalized in the embryo during germ band extension (Seydoux and Dunn, 1997). We note that unlike *D. melanogaster,* both *O. fasciatus* and *G. bimaculatus* appear to specify primordial germ cells through inductive signals rather than germ plasm (Donoughe et al., 2014; Ewen-Campen et al., 2013; Nakamura and Extavour, 2016) , and that transcriptional activity is required to respond to inductive signaling. In *M. musculus*, another animal in which primordial germ cells are specified through inductive signaling (Lopes et al., 2007, 2004; Ying and Zhao, 2001), primordial germ cells are transcriptionally quiescent for only a brief period during their migration (Seki et al., 2007). It is possible that a more thorough analysis in future studies of H5 antibody stains may reveal a similar period of germ cell transcriptional quiescence in either *O. fasciatus* or *G. bimaculatus*.

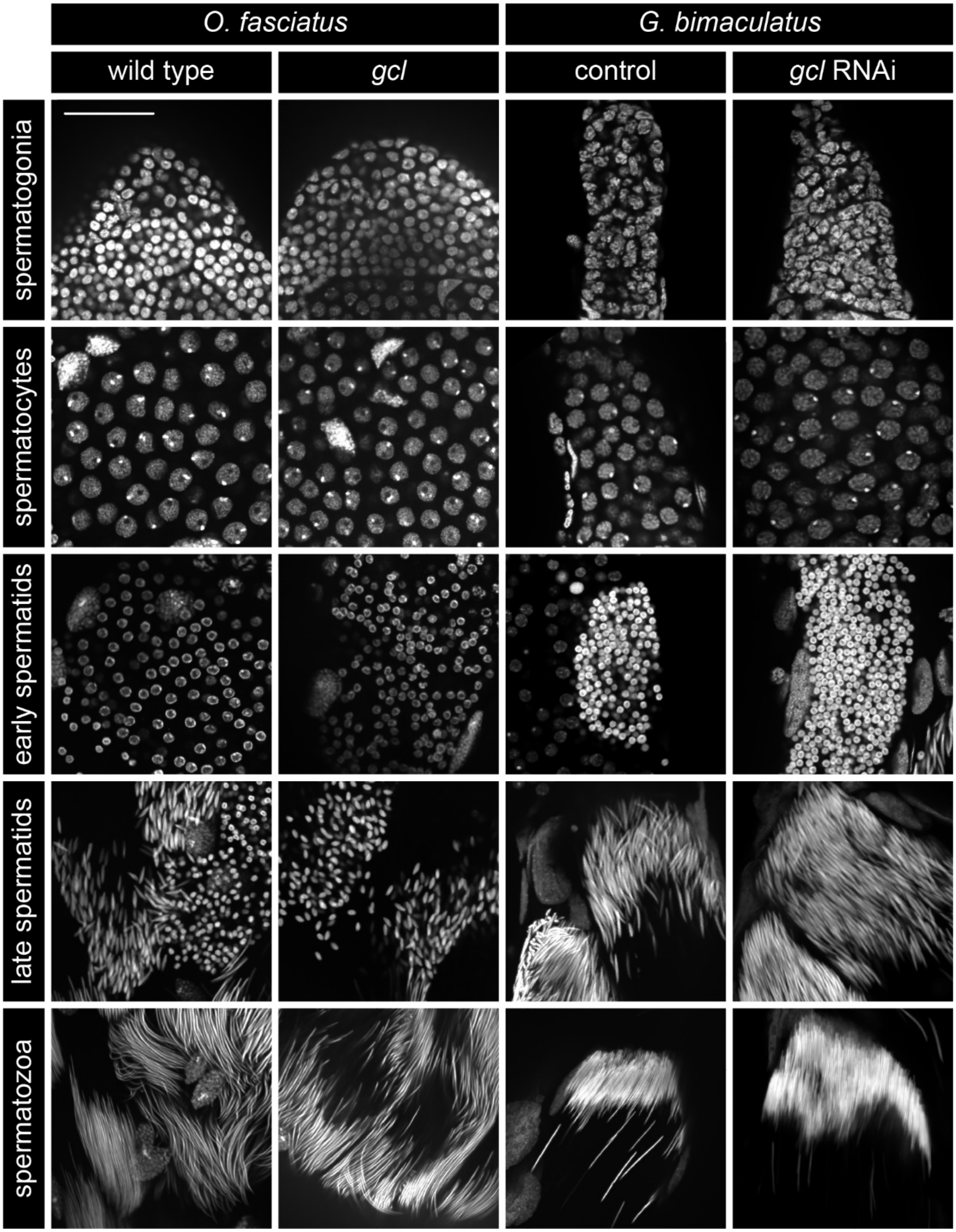


**Supplemental Figure 4. Lack of *gcl* activity does not affect the progression of adult spermatogenesis in either *O. fasciatus* or *G. bimaculatus*.** Representative images of Hoechst-stained testes showing different stages of spermatogenesis in both species under both wild type and *gcl* loss of function conditions. Scale bar = 50 μm.

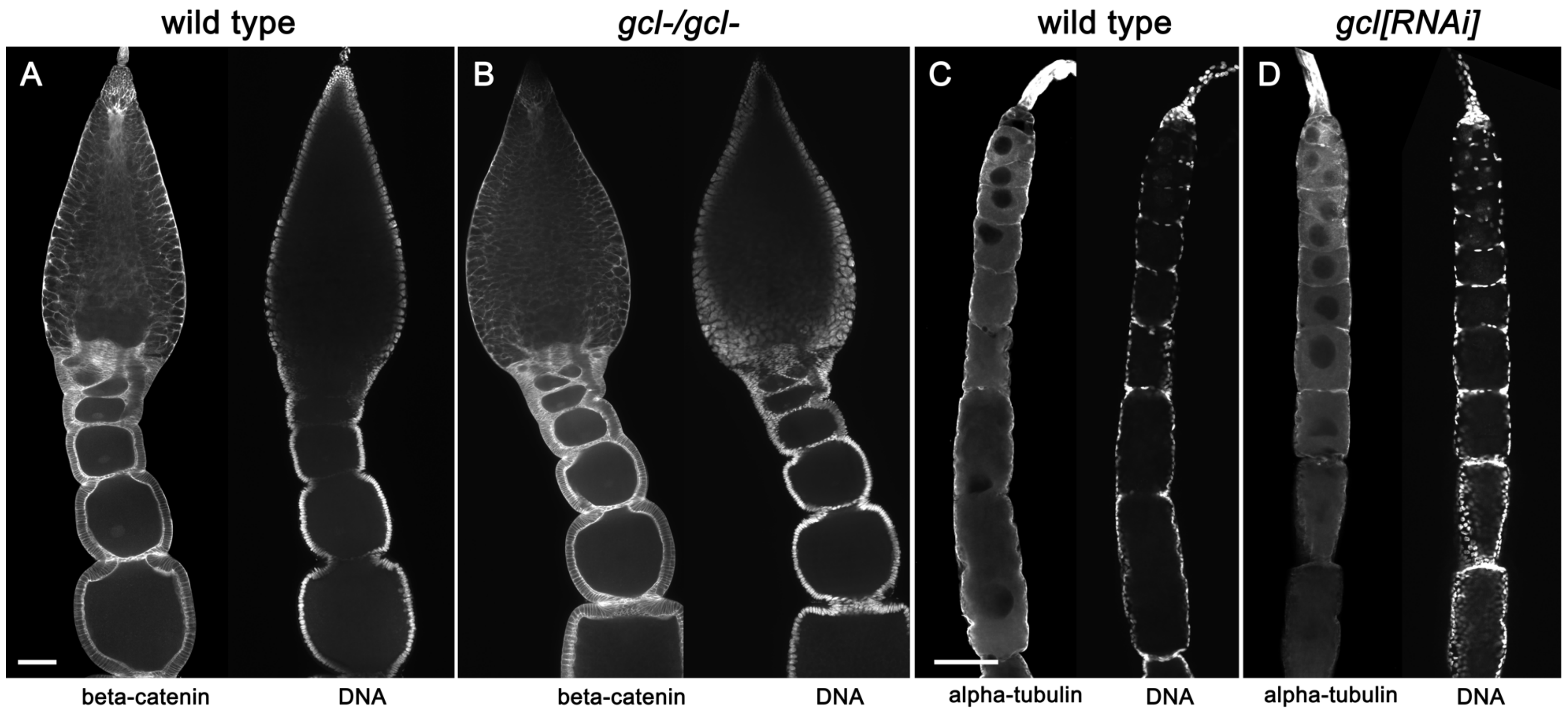


**Supplemental Figure 5. Lack of *gcl* activity does not affect adult oogenesis in either *O. fasciatus* or *G. bimaculatus*.** Representative images of single 1 µm optical sections of ovarioles under both wild type ((A) *O. fasciatus*; (C) *G. bimaculatus*) and (B) *gcl* knockout (*O. fasciatus*) or (D) knockdown (*G. bimaculatus*) conditions. Scale bars = 100 µm in (A) (applies also to (B)) and (C) (applies also to (D)).

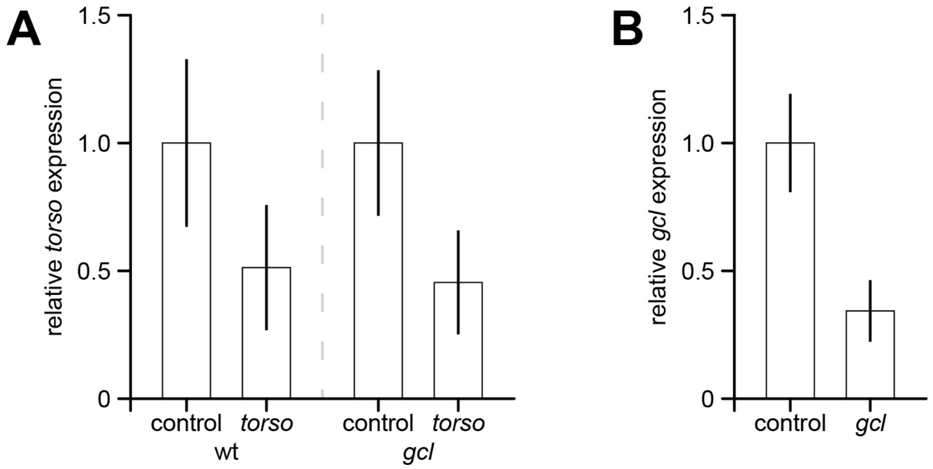


**Supplemental Figure 6. qPCR validation of RNAi experiments.**

(**A**) Efficacy of *torso* RNAi in *O. fasciatus* embryos. $C_q$ values were normalized against *Of-gapdh*, averaged across three biological replicates for each condition, and normalized to the control average. *torso* RNAi applied within four hours of egg-laying reduces detectable transcript levels to ~50% of control levels at 24 hours after injection in both wild type and *gcl* mutant backgrounds.

(**B**) Efficacy of *gcl* RNAi in *G. bimaculatus* embryos. $C_q$ values were normalized against *Gb-tubulin*, averaged across three biological replicates for each condition, and normalized to the control average. *gcl* RNAi applied within 4 hours of egg-laying reduces detectable transcript levels to ~35% of control levels in stage 8 embryos, four days after egg-laying.

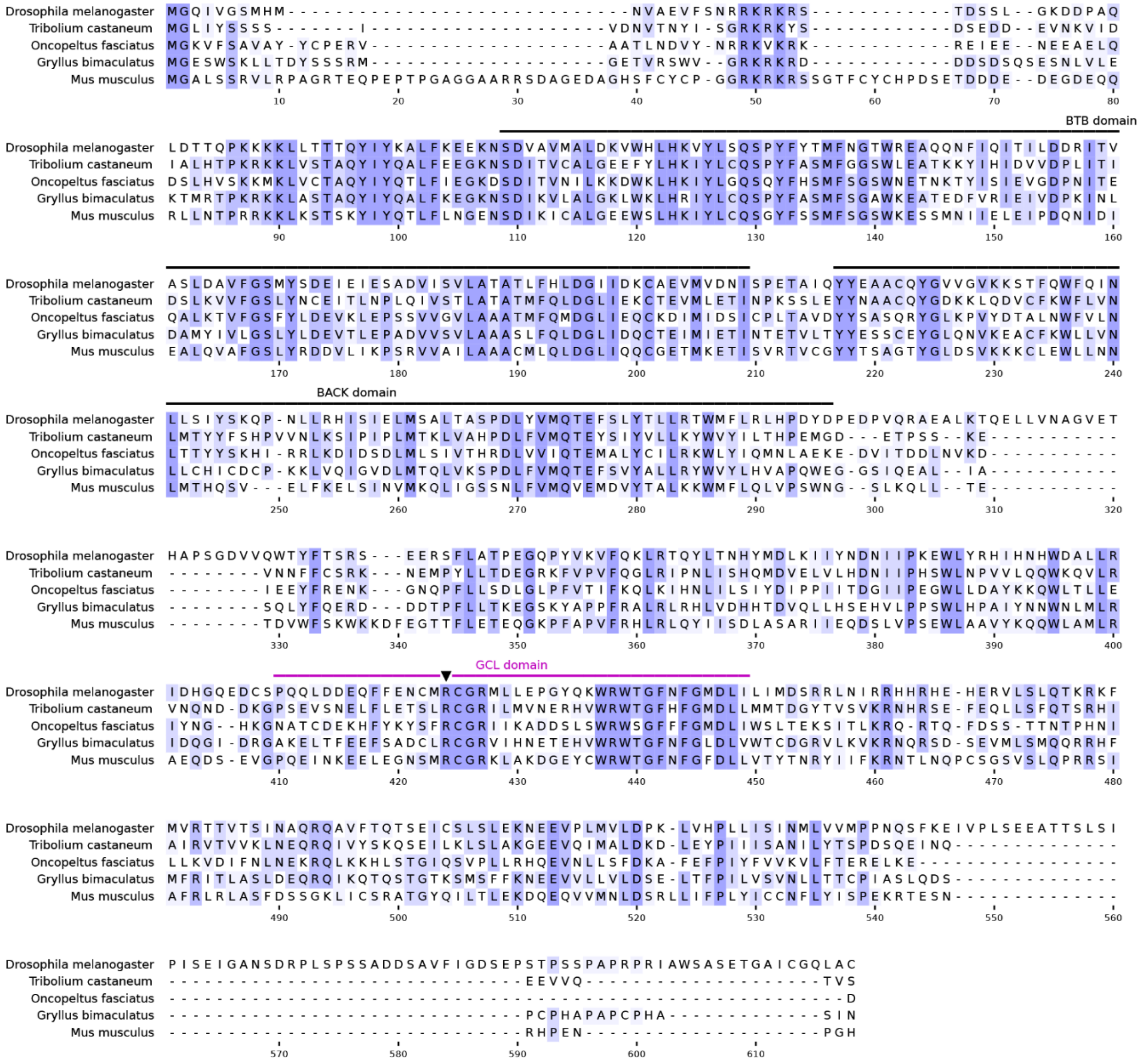


**Supplemental Figure 7. MUSCLE alignment of select *gcl* homologs.** Darker shading represents greater degree of conservation at each position. The conserved GCL domain identified by Pae and colleagues (Pae et al., 2017) is marked in magenta. In that study, the role of the GCL domain was directly tested by mutating the residue marked by the arrowhead (R377).

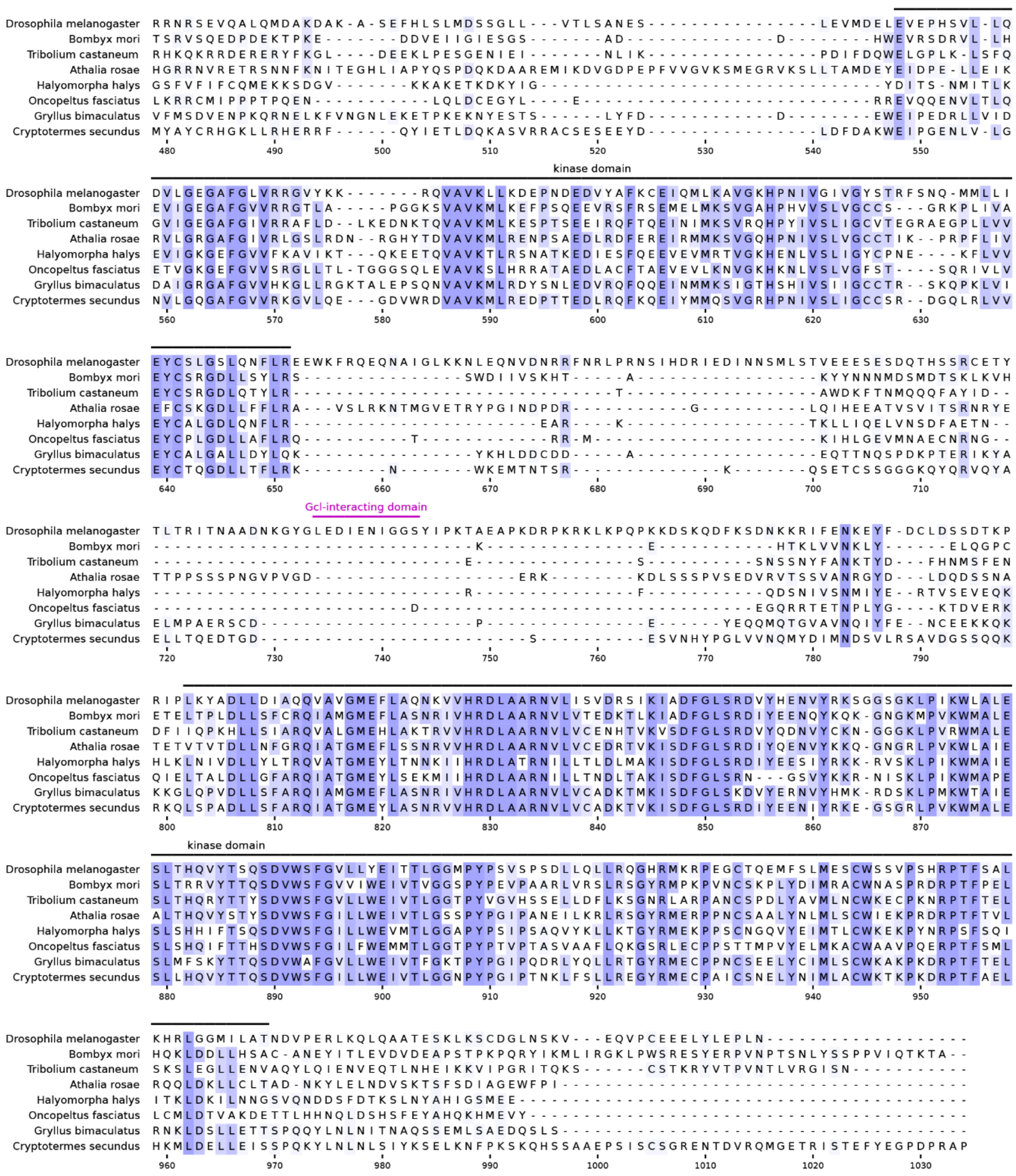


**Supplemental Figure 8. MUSCLE alignment of the intracellular domain of select insect *torso* homologs.** Darker shading represents greater degree of conservation at each position. The Gcl-interacting motif identified by Pae and colleagues (Pae et al., 2017) is marked in magenta. Note that this motif is not conserved in other insects; the surrounding domain is much longer in *D. melanogaster* compared with other homologs.

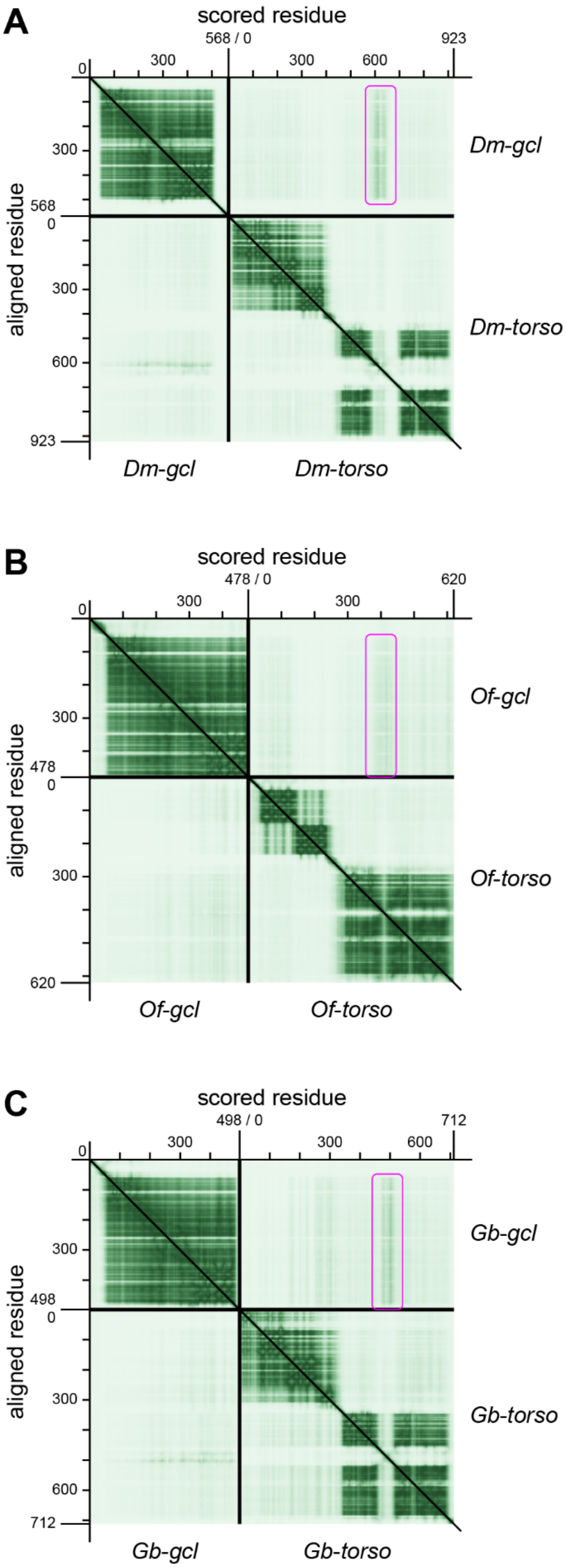


**Supplemental Figure 9. AlphaFold3 predicts potential interactions between Gcl and Torso homologs.** Predicted alignment error plots are shown for AlphaFold3 structure predictions of Gcl and Torso homologs from *D. melanogaster* (**A**), *O. fasciatus* (**B**) and *G. bimaculatus* (**C**). Darker color represents higher confidence (lower predicted error) in the relative positions of the scored and aligned residues. Overall confidence in the structure of the Gcl–Torso complex is low for all species. However, in all species, there is a region of Torso protein (marked in magenta) for which its predicted alignment error with Gcl is lower, suggesting a potential interaction between the two proteins through this region of Torso.

**Supplemental Table 1. Primers used.**

| gene | direction | primer sequence | expected length |
|---|---|---|---|
| full-length sequence | | | |
| *Of-gcl* | forward | ATGGGTAAAGTATTCAGTGCAGTTGCT | 1437 bp |
| | reverse | TTAGTCTTCTTTTAATTCTCTCTCTGTAAATAG | |
| *Gb-gcl* | forward | CGTCGATGAGCTAGTTGAGAGCTGA | 1804 bp |
| | reverse | AGATGTTCAAGCATAGCCGCCCAAG | |
| *in situ* probe and dsRNA | | | |
| *Of-gcl* | forward | GCAGTTGCTTATTACTGCCCAGAGCGT | 441 bp |
| | reverse | TGCAGCAGCCAGCACACCAACAACA | |
| *Of-torso* | forward | TCGAGGTCGCCGTCAAATCCCT | 665 bp |
| | reverse | GGCAACAGACGCTGTCGGAACA | |
| *Gb-gcl* | forward | TGTCCGCTCGTGGGTCGGCAGAAAA | 413 bp |
| | reverse | ACAGAGAGGCTGCAGCAAGCACTGA | |
| qPCR | | | |
| *Of-gcl* | forward | TCCAGAAGGTTGGTTGCTCGATGCCT | 257 bp |
| | reverse | TGCTGCTGTCAAACTGTGTACGCTGCC | |
| *Of-torso* | forward | TGAGCATGGTGGGCGATTGCTG | 288 bp |
| | reverse | AGGGATTTGACGGCGACCTCGA | |
| *Gb-gcl* | forward | GCACCACCATTTCGTGCACTTCGCC | 258 bp |
| | reverse | ACCTGTCCATCGCCACACATGTTCT | |
| *Of-gapdh* | forward | TGGCAAACTAACTGGCATGGCA | 171 bp |
| | reverse | ACAACCTGGTCATCAGTGTACGCCA | |
| *Gb-β-tubulin* | forward | TGGACTCCGTCCGGTCAGGC | 166 bp |
| | reverse | TCGCAGCTCTCGGCCTCCTT | |
| RACE | | | |
| *Of-torso* | forward 1 (3' RACE) | ACTGGAGGAGGCTCCCAACTCGAGGTCGCC | |
| | forward 2 (3' RACE) | ACAGGAGGGCCACGGCAGAGGACTTGGCTT | |
| | reverse 1 (5' RACE) | TGGAGAAAGGCGGCAACAGACGCTGTCGGA | |
| | reverse 2 (5' RACE) | CGCTGGGAGGTAGAGAAGCCGACCAGGGAG | |
| *Of-fgfr* | forward 1 (3' RACE) | AACAAGGGCATACAGATGCAGAAATGACTG | |
| | forward 2 (3' RACE) | GCATCGGCCATCATCTGGTTATGAACCTGC | |
| | reverse 1 (5' RACE) | TATTCCTTGCTGCCAGGTCTCGATGTAAAC | |
| | reverse 2 (5' RACE) | TGACAGGTATTCCATGCCTCTAGCAACTTG | |

**Supplemental Table 2. Staging table for wild type *O. fasciatus* embryogenesis at 28°C.**

| stage | hours ael (28°C) | stage description | germ cell development |
|---|---|---|---|
| 1 | | Freshly laid egg. | |
| 2 | | Uniform blastoderm. | |
| 3 | 26–28 | An indentation appears at the posterior end of the egg, called the posterior pit. | Primordial germ cells are first observed just inside the posterior pit, but we cannot currently perform antibody stains at this stage. |
| 4 | 28–32 | The germ band has invaginated into the posterior pit and extends toward the anterior. The orientation of the germ band is reversed with respect to the egg, *i.e.*, the posterior end of the germ band is closer to the anterior of the egg. | Primordial germ cells remain in a cluster at the posterior tip of the germ band. The Vasa antibody stain is inconsistent at this stage. |
| 5 | 32–36 | The thorax region is wider than the abdomen region (mediolaterally), but limb buds are not yet distinct. Segments have been specified up to A5–A7 but are not yet distinguishable without staining for a segmentation marker. | Germ cells are located in the bulbous part of the germ band, primarily in A5–A6, but with some in A4 and A7. |
| 6 | 36–44 | Limb buds are now distinct semicircles. Segments have been specified up to A7–A9 but are not yet distinguishable without staining for a segmentation marker. | Germ cells are located in a single cluster primarily in A4–A6, with some in A7. Segments in which germ cells are located are narrower in the medial–lateral axis compared with the posterior tip. |
| 7 | 42–50 | Segments start to become visibly distinct in the anterior abdomen around the time A9 is specified. Posterior tip is still round. Neural groove has formed at most up to A3. | Germ cells separate into segmental clusters. When there are many germ cells, they may still appear as one large medial cluster, especially as one spanning both segments A4 and A5. |
| 8 | 48–54 | Legs and gnathal appendages are triangular rather than semicircular lobes. A11 starts to thicken but the posterior tip is still mostly round. | Germ cells split into bilateral clusters as the neural groove forms in each segment. In this stage, germ cells are located right at the edges of the neural groove. When there are many germ cells in a particular segment, they may still appear to be one large medial cluster in that segment. |
| 9 | 52–64 | Neural groove has formed up to A8–A9. "Coelomic cavities" are visible as pits in all abdominal segments up to A9, though they may sometimes be obscured by germ cells, especially in A4–A6. Posterior end has folded over to form the proctodeum. Lateral | Germ cells have moved laterally away from the neural groove and clusters usually appear rounder. |

| | | | |
|---|---|---|---|
| | | margins of the germ band are thickening in the gnathal and thoracic segments. | |
| 10 | 60–64 | Lateral margins of the germ band are thickening in the abdomen. Ganglia appear as two parallel rectangles in each segment, separated by the median cord along the midline. “Coelomic cavities” are no longer distinct. | Germ cell clusters lengthen along the AP axis and narrow along the medial–lateral axis. Adjacent clusters may fuse, but a segmental grouping is still visible. |
| 11 | 64–68 | Median cord clusters appear at segmental boundaries, and the ganglionic masses are now slightly crooked. | Germ cells are lined up in the genital ridges on each side of the embryo, parallel to the AP axis. |
| 12 | 68–72 | Abdominal cross commissures have formed. With a nuclear stain, ganglia appear as pairs of half circles facing away from each other. | Germ cells remain in the genital ridges. |
| 13 | 72–76 | The lateral cords move dorsolaterally and become parallel with the median cord again. Gonadal follicle primordia become visible in the nuclear channel. | Germ cells remain in the genital ridges. |
| 14 | 76– | The embryo undergoes katatrepsis and begins dorsal closure. This stage will likely be subdivided further. | Germ cells start to migrate laterally into gonadal follicle primordia. |

**Supplemental Table 3. Quantification of germ cell development in wild type and *gcl* mutant *O. fasciatus* embryos.** All *p*-values were calculated using a non-parametric bootstrap test.

| | total germ cells per embryo | | | | | | proportion of PH3+ germ cells per embryo | | | | |
|---|---|---|---|---|---|---|---|---|---|---|---|
| | wild type | | *gcl* | | | | wild type | | *gcl* | | |
| stage | mean | *n* | mean | *n* | ratio | *p*-value | mean | *n* | mean | *n* | *p*-value |
| 3 | 102.00 | 14 | 68.13 | 8 | 0.668 | <0.001 | 0.0126 | 14 | 0.0110 | 8 | 0.4221 |
| 4 | 108.00 | 22 | 56.55 | 20 | 0.524 | <0.001 | 0.0181 | 22 | 0.0124 | 20 | 0.0953 |
| 5 | 115.69 | 54 | 77.58 | 36 | 0.671 | <0.001 | 0.0221 | 49 | 0.0279 | 36 | 0.3386 |
| 6 | 124.71 | 52 | 65.70 | 20 | 0.527 | <0.001 | 0.0250 | 47 | 0.0236 | 20 | 0.0382 |
| 7 | 133.54 | 59 | 86.19 | 26 | 0.645 | <0.001 | 0.0137 | 46 | 0.0143 | 26 | 0.2572 |
| 8 | 118.57 | 28 | 75.70 | 20 | 0.638 | <0.001 | 0.0108 | 22 | 0.0132 | 20 | 0.0745 |
| 9 | 122.19 | 65 | 96.36 | 14 | 0.789 | 0.002 | 0.0081 | 43 | 0.0067 | 14 | 0.0164 |
| 10 | 130.24 | 21 | 79.77 | 13 | 0.612 | <0.001 | 0.0040 | 17 | 0.0058 | 13 | 0.4213 |
| 11 | 113.36 | 31 | 80.52 | 25 | 0.710 | 0.001 | 0.0040 | 30 | 0.0045 | 25 | 0.4566 |
| 12 | 91.04 | 26 | 70.29 | 17 | 0.772 | 0.017 | 0.0030 | 23 | 0.0042 | 17 | 0.2247 |
| 13 | 89.95 | 39 | 68.63 | 35 | 0.763 | <0.001 | 0.0014 | 26 | 0.0012 | 35 | 0.1769 |
| 14 | 101.67 | 33 | 69.03 | 30 | 0.679 | <0.001 | 0.0015 | 33 | 0.0005 | 30 | 0.0605 |
